# An updated *C. elegans* nuclear body muscle transcriptome for studies in muscle formation and function

**DOI:** 10.1101/2022.04.12.488068

**Authors:** Anna L. Schorr, Alejandro Felix Mejia, Martina Y. Miranda, Marco Mangone

## Abstract

The body muscle is an important tissue used in organisms for proper viability and locomotion. Although this tissue is generally well studied and characterized, and many pathways have been elucidated throughout the years, we still lack a comprehensive understanding of its transcriptome and how it controls muscle development and function. Here, we have updated a nuclear FACS sorting-based methodology to isolate and sequence a high-quality muscle transcriptome from *C. elegans* mixed-stage animals. We have identified 2,848 muscle-specific protein-coding genes, including 78 transcription factors and 206 protein-coding genes containing an RNA binding domain. We studied their interaction network, performed a detailed promoter analysis, and identified novel muscle-specific *cis*-acting elements. We have also identified 16 high-quality muscle-specific miRNAs, studied their function *in vivo* using fluorochrome-based analyses, and developed a high-quality *C. elegans* miRNA Interactome incorporating other muscle-specific datasets produced by our lab and others.

Our study expands our understanding of how muscle tissue functions in *C. elegans* and in turn, provide results that can in the future be applied to humans to study muscular-related diseases.

## INTRODUCTION

The body muscle is an important tissue used in organisms for proper viability and locomotion. Muscle tissue’s structure, shape, and size change depending on its function and location. In humans, there are three main types of muscles: skeletal, smooth, and cardiac muscles, each responsible for a different form of contraction. Smooth and cardiac tissue contract involuntarily and are mainly located on the surface of organs. In contrast, skeletal muscles contract voluntarily.

The contractile unit of the muscle is the sarcomere. It is composed of at least thirty different proteins, of which the most abundant are myosin and actin, and is ultimately responsible for the contraction reaction leading to movement.

The model organism *C. elegans* is ideal for studying muscle development and morphogenesis due to its well-characterized transcriptome, fully mapped cell lineage, and the availability of several methods for genetic engineering.

The basic contractile unit of the muscle is the sarcomere, which is strongly conserved in its structure and function from *C. elegans* to mammals (WormBook). *C. elegans* sarcomeres are composed of myosin-containing thick filaments and actin-containing thin filaments, which are responsible for the contraction reaction leading to movement through the myosin heads pulling on the actin filaments (WormBook). A notable difference between the worm and vertebrates is the absence of the costamere and Z-disc, which in vertebrates anchors the actin filaments to the extracellular matrix. However, worms contain a structure called the dense body, which performs the same function [1]. In addition, over 200 proteins have been identified in *C. elegans* to be essential for the assembly, maintenance, and function of the sarcomere, many of which have human orthologs [2]. *C. elegans* is also a popular choice for studying disorders affecting humans, including muscular dystrophy [3–5], since many human disease genes, when introduced in worms phenocopy the symptoms [6–9].

*C. elegans* possess two large muscles: the pharynx and the body muscle. The pharynx is localized to the front of the animal and is responsible for food intake and the physical crushing [10]. It is composed of eight layers of non-striated muscles, surrounded by epithelial and neural tissue. The body muscle tissue comprises 95 striated muscle cells localized throughout the animal’s body and is functionally equivalent to the vertebrate skeletal muscles [11]. The contractile unit of the muscle is the sarcomere. It is composed of at least 30 different proteins, of which the most abundant are myosin and actin, and is ultimately responsible for the contraction reaction leading to movement. The sarcomere is highly conserved between nematodes and vertebrates and is composed of repeated units of a complex mesh of thin (I-band) and thick (A-band) filaments. The thin filaments are mainly composed of actin and anchor each sarcomere to the dense bodies and stabilize the entire structure. Conversely, the thick filaments are mostly made of myosin and are attached to the M-line and mediate tension generated during muscle contraction.

To properly develop and maintain functional body muscle tissue, gene expression must be strictly regulated. Our group and others in the past finely characterized the transcriptome of the *C. elegans* body muscle in the adult stage [12, 13] and during the embryonic development [14] and identified important regulatory mechanisms used by muscle cells to maintain their homeostasis.

In addition to the identification of important genes in this tissue, our group and others recently identified that tissue-specific alternative polyadenylation (APA), a poorly characterized mechanism that produces genes with different 3’untranslated region (3’UTR) isoforms, is an important regulatory mechanism used by cells to evade the negative regulation effected by microRNAs (miRNAs) [12, 13]. Still, the complexity and depth of this form of regulation are currently unknown. Regulation of gene expression also depends on miRNAs, which are short ncRNAs that target regulatory elements in 3’UTRs and repress gene expression in the cytoplasm [15, 16].

Since their discovery, several groups, including our lab, have shown that miRNAs influence gene regulation considerably [17]. The canonical miRNA biogenesis pathway begins with the transcription of pri-miRNAs in the nucleus, which fold into a hairpin structure [17]. The microprocessor complex cleaves the pri-miRNAs into a shorter stemloop structure, the pre-miRNAs, which are then exported to the cytoplasm [17]. Once in the cytoplasm, the pre-miRNAs are cleaved into mature miRNAs, which associate with an Argonaute protein as part of the RNA-induced silencing complex (RISC) [17–19]. When loaded into the RISC, miRNAs bind to complementary target sequences within the 3’UTRs of genes to be repressed [17]. This implies that the 3’UTRs and their isoforms are integral for miRNA-based regulation.

Several strategies have been developed to sequence tissue-specific transcriptomes and miRNAs in *C. elegans*. These methodologies utilize either fluorescence-activated cell sorting (FACS) [14, 20], immunoprecipitation [12, 13, 21], or RNA modification techniques [22]. Each of these approaches has unique advantages but also presents several caveats. Methodologies utilizing FACS allow for high purity and stringency when isolating RNA from cells or nuclei. These approaches rely on tissue-specific promoters to drive fluorochromes expression in cells or nuclei. After isolation, the fluorescent cells or nuclei may be readily sorted using FACS, then either sequenced [20] or processed using microarrays [14].

Immunoprecipitation and antibody-based approaches have also been widely used to isolate the transcriptomes of specific tissues. This approach provides considerably high yields of RNA compared to FACS-based methods and is generally much more cost-efficient.

Brosnan *et al*. (2021) used an epitope labeled Argonaute protein expressed in specific *C. elegans* tissues to immunoprecipitate tissue-specific miRNAs populations from the body muscle, intestine, and neurons [21].

Alberti *et al*. (2018) developed a methodology named “microRNAome by methylation-dependent sequencing (Mime-seq)” to identify tissue-specific miRNA populations from complex animals at the level of single-cell specificity [22]. The Mime-seq approach expresses a transgenic methyltransferase (Ath-HEN1) from *Arabidopsis thaliana* in an animal system, allowing the labeling of small RNAs. Mime-seq has been utilized in *C. elegans* and *Drosophila* and improved tissue-specific miRNA identification compared to previous immunoprecipitation-based methods [22].

Our group also used an immunoprecipitation-based approach to identify transcripts targeted by miRNAs within the *C. elegans* intestine, and body muscle [23] and observed transcripts were targeted differentially between the two tissues. While identifying the miRNA targets provided valuable information regarding tissue-specific gene regulation, a major limitation of this study was that we could not identify the specific miRNAs involved.

Here, we describe a novel nuclear FACS sorting-based approach named “Nuc-Seq,” to allow the isolation and sequencing of high-quality muscle transcriptomes and miRNA populations from *C. elegans* mixed-stage animals. Using this method, we have identified 2,848 muscle-specific protein-coding genes, studied their interaction network, performed a detailed promoter analysis, and identified novel muscle-specific *cis*-acting elements. Using fluorochrome-based analyses *in vivo*, we developed a high-quality muscle miRNA interactome, which incorporates other muscle-specific datasets produced by our lab and others.

## RESULTS

### Nuc-Seq: an updated approach to identify muscle-specific transcriptome

We wanted to improve the annotation of the body muscle transcriptome by developing a nuclear FACS-based strategy to isolate and sequence transcriptomes from *C. elegans* body muscle nuclei. We named this approach ‘Nuc-Seq.’ Our final goal was to identify the body muscle transcriptome and its miRNA population expressed explicitly in this tissue.

To test the feasibility and optimize this approach, we first performed experiments using the *C. elegans* strain BN452, which ubiquitously expresses the mCherry fluorochrome fused to the histone H2B ortholog gene *his-58* [24] (**Main Figure 1**). We fractionated this worm strain using mechanical stress and separated two fractions: one cytoplasmic and one enriched with mCherry fluorescent nuclei. (**Main Figure 1A-B and Materials and Methods**). Using a Western blot approach, we successfully tested these two fractions for the presence of known cytoplasmic and nuclear proteins (**Main Figure 1C**). Importantly, these nuclei are stable over time with minimal degradation after several hours (**Supplemental Figure S1**). We then performed the FACS sorting step and successfully isolated a large pool of mCherry-positive nuclei, with over 34% of all the events being mCherry-positive (**Main Figure 1D**). We named this updated approach Nuclear Sequencing (Nuc-Seq).

**Main Figure 1:**
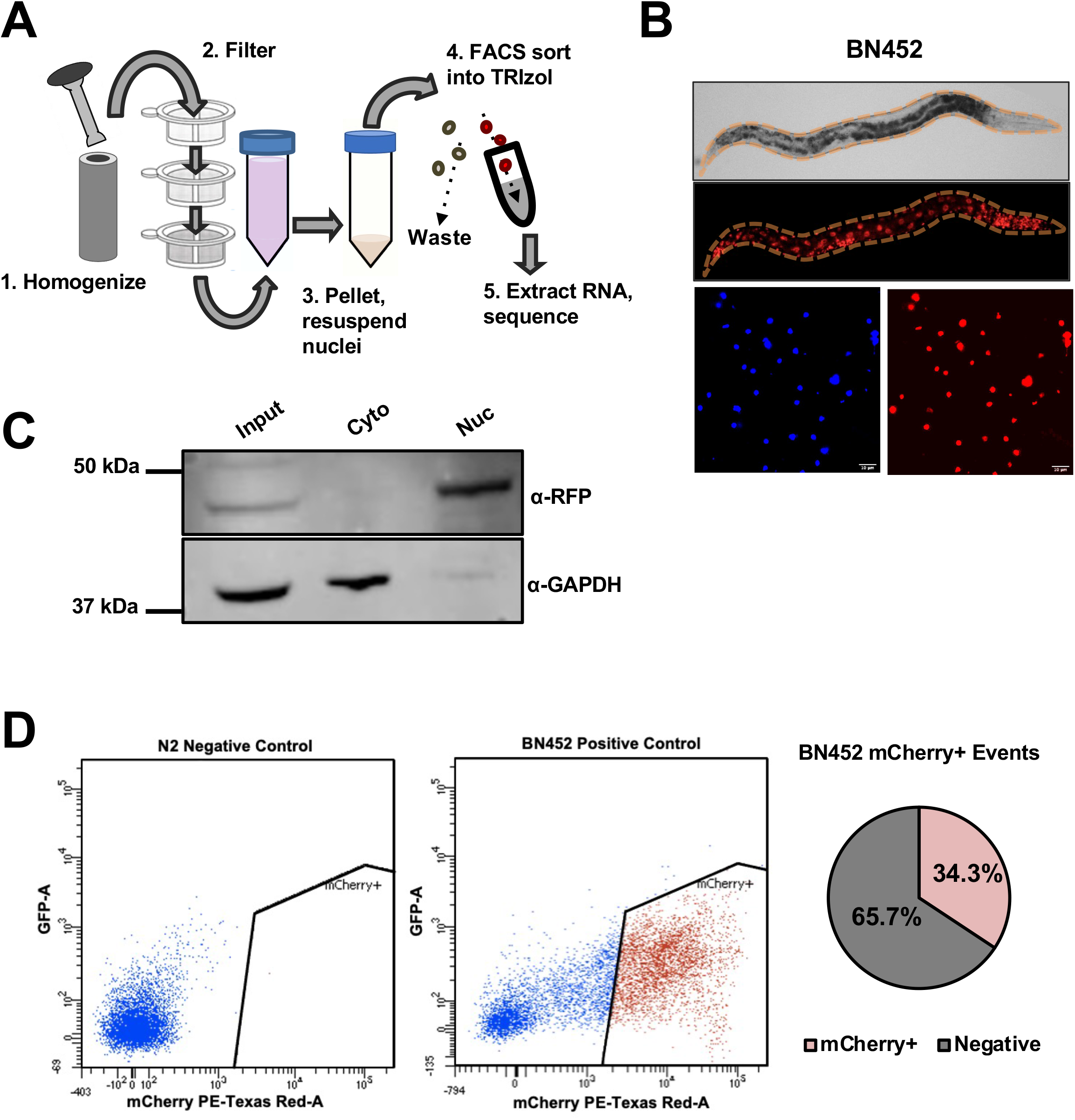
Nuc-Seq schematics. A) Transgenic worms expressing mCherry fluorochrome fused to histone H2B (*his-58*) are (1) homogenized and (2) then subjected to several rounds of mechanical filtration. (3) The resultant nuclei are then collected and (4) subjected to FACS sorting. After the sorting step, (5) the mCherry-positive nuclei were sorted directly into TRIzol, and the recovered RNA was sequenced. B) Top panel: The strain BN452 ubiquitously expresses the mCherry:: *his-58* transgene. Bottom Panel: mCherry positive nuclei recovered after Step 2 described in Panel A. (Red: mCherry, Blue: DAPI). C) Western blot of nuclear and cytoplasmic fractions from BN452 worms blotted with α-RFP and α-GAPDH antibodies. D) Left Panel: nuclear FACS profile from N2 and BN452 nuclei. The BN452 sample is a positive control that represents all nuclei. The red dots show the sorted mCherry-positive population. Right Panel: Pie chart showing the percentage of BN452 sorted mCherry-positive nuclei.

With these results in our hands, we decided to move forward and limit the expression of the *mCherry::his-58* cassette in the body muscle tissue. We cloned the *his-58* gene, fused it to the mCherry fluorochrome, and then forced its expression only in the body muscle tissue using the promoter region of the *C. elegans* ortholog of the myosin heavy chain gene (*myo-3*) (**Main Figure 2A upper panel**). As expected, the resultant transgenic worm restricts the expression of the mCherry fluorochrome in the body muscle nuclei (**Main Figure 2A lower panel**), with fewer fluorescent nuclei compared to the experiments in the BN452 strain (**Main Figure 2A**). We then performed the FACS sorting step (**Main Figure 2B**). Although the number in sorted nuclei is significantly less than those obtained with the positive control BN452, we could still isolate a large population of mCherry-positive nuclei across from each of our replicates, which we then extracted and sequenced RNA from.

**Main Figure 2:**
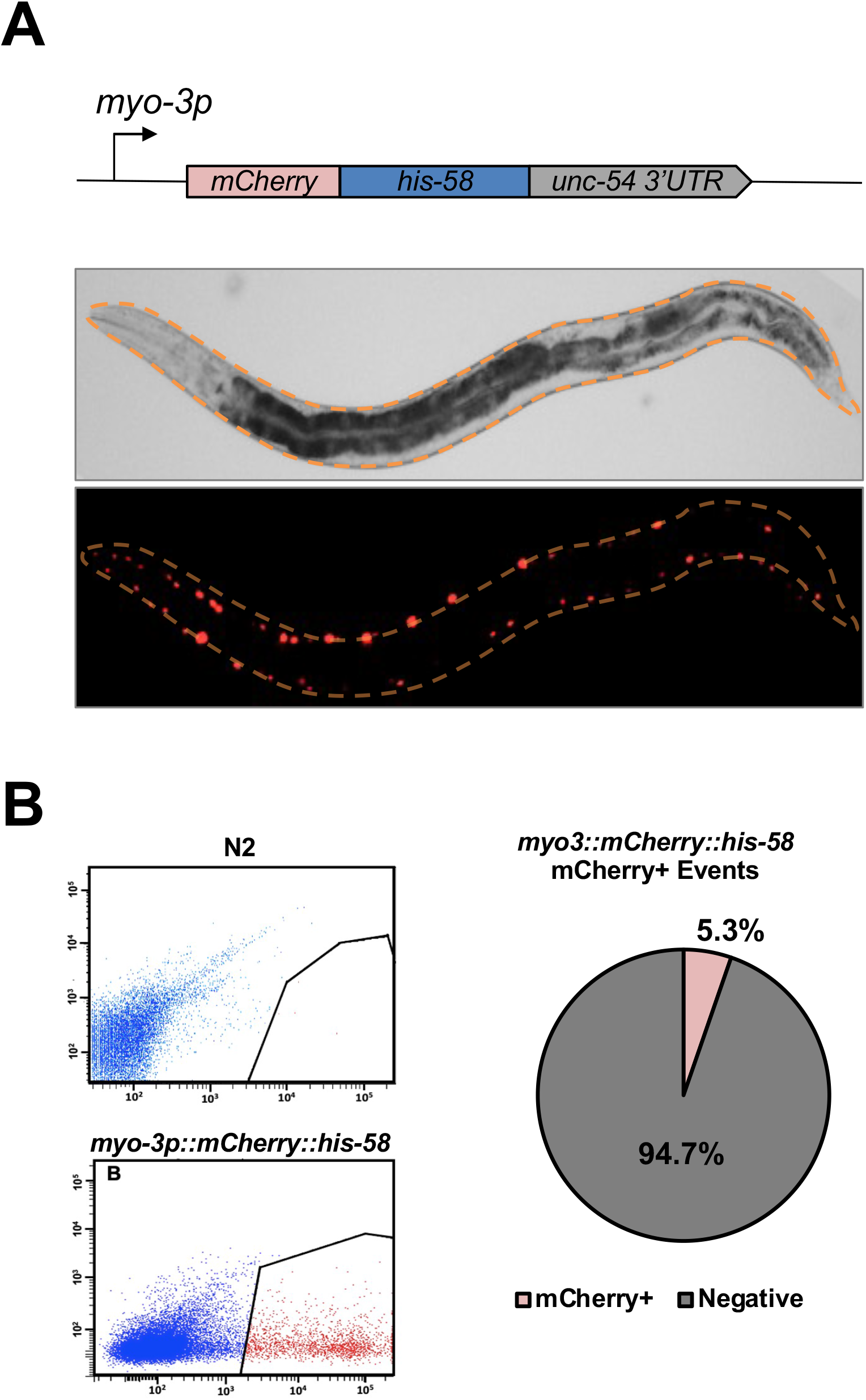
FACS sorting of body muscle nuclei: A: Top Panel: Schematics of the construct used to isolate the body muscle transcriptome. The body muscle-specific promoter *myo-3* drives the expression of the mCherry:::*his-58* cassette. Bottom Panel: Bright field and fluorescent images of the resultant transgenic worm strain. B) nuclear FACS profile from N2 and *myo3p::mCherry:::his-58::unc-54 3’UTR* nuclei. The red dots show the sorted mCherry-positive population. Right Panel: Pie chart showing the percentage of sorted mCherry-positive and body muscle-specific nuclei.

### The identification of the C. elegans body muscle transcriptome

We performed three sequencing reactions in duplicates (technical triplicates), two from our body muscle-expressing nuclei (body muscle enriched), and one from nuclei isolated from BN452 (negative controls) (a total of six samples). We obtained approximately 55M mappable reads for each sample (including our technical replicates). For most of the samples, we could map more than 97% of the total reads (**Supplemental Figure S2A**). The results obtained with our biological replicates correlate well (**Supplemental Figure S2B-C**).

We have identified 2,849 protein-coding genes in the *C. elegans* body muscle tissue, corresponding to ~14% of all *C. elegans* protein-coding genes (20,362 protein-coding genes; WS250) (**Main Figure 3 and Supplemental Table S1**). Using comparative data from Kim et al. [25], we identified human orthologs for 1,872 worm genes (66%).

**Main Figure 3:**
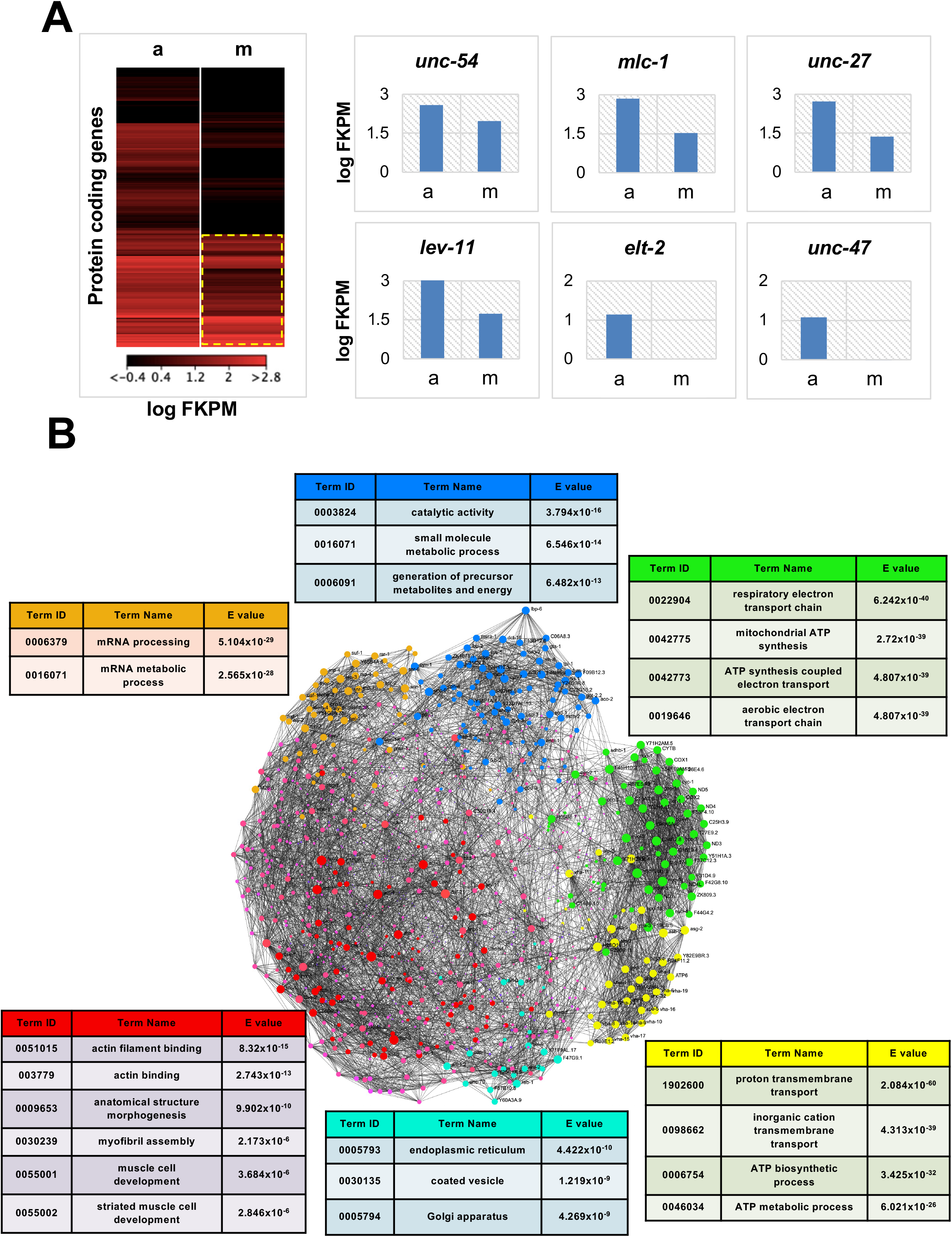
The *C. elegans* body muscle transcriptome. A) Left Panel: Heat map showing the body muscle expression levels log(fpkm) of the 20,362 protein-coding genes in WS250. (a) BN452 positive control, (m) muscle transcriptome. The body muscle-specific transcriptome is highlighted with a dashed yellow line. Right Panel: several examples of expression levels in genes identified in the (a) BN452 (ubiquitously expresses GFP and mCherry in all nuclei; positive control) and (m) body muscle-specific nuclear fractions. The vertical axes mark expression levels as the log(FPKM). The body muscle-specific genes *unc-54, mlc-1, unc-27*, and *lev-11* were all detected in our study, while the intestine-specific *elt-2* and the GABA neuron-specific *unc-47* were not present. (a: median log(fpkm) of the ‘BN452 all nuclei’ dataset; m: median fpkm values of set1 and set2 muscle datasets). B) The *C. elegans* body muscle Interactome. We clustered all 2,849 protein-coding genes identified in this study using previously published protein-protein interaction data. Each gene is shown as a pink dot. Within this network, we identified six smaller, highly interconnected subnetworks shown in different colors. The genes in each subnetwork were independently subjected to GO term enrichment analyses shown with colored panels above each respective subnetwork.

While some of our top hits have not been characterized yet, as expected, we detected many known components of the sarcomere, such as many of the proteins that form the thick filaments (myosin heavy chain genes, *myo-3, unc-54*) and myosin light chain (*mlc-1*), and sarcomere organization and structural factors (*unc-15, unc-45*), as well as those that form the thin filaments, including several actin genes (*act-1, act-3, act-4*) and structural factors (*unc-60, unc-73*) (**Main Figure 3A and Supplemental Figure S3)**. We also detect orthologs of the tropomyosin gene *lev-11*, which in resting sarcomere blocks myosin binding to actin, and other notable genes previously known to be in the *C. elegans* body muscle tissue (**Main Figure 3A and Supplemental Figure S3)**.

Our dataset identified 78 transcription factors, including *mdl-1, skn-1, xbp-1, mxl-3*, all previously known to be expressed in the body muscle tissue, and 207 non-ribosomal proteins containing an RNA binding domain **Supplemental Table S1**. Within this group, we identified K08D12.3, an ortholog of the human ZNF9, which is mutated in myotonic dystrophy type 2, *mbl-1*, a MUSCLEBLIND-type of mRNA splicing regulator required for muscle dense body organization, locomotion, and vulval morphogenesis, and *etr-1*, an ortholog of the human CUG-binding protein CUGBP1, which is required for embryonic muscle development and has been implicated in myotonic dystrophy **Supplemental Table S1**.

Next, we studied the interaction network of the identified C. *elegans* muscle genes. We used the STRING database [26] for this analysis, which contains known *C. elegans* predicted and functional protein-protein interactions. Using our top 2,000 genes, we produced a large network containing 1,995 nodes and 31,449 edges (**Main Figure 3B)**. The genes are highly interconnected (**Main Figure 3B),** producing six large distinguished sub-networks involved in body muscle formation, maintenance, protein production, and energy storage (**Main Figure 3B)**. A GO term analysis shows functional enrichments for actin filament binding, skeletal muscle myosin filament assembly, mitochondrial functions, and ATP biosynthesis: all processes functioning in the body muscle tissue (**Main Figure 3B)**. 71% of all genes identified in this study are protein-coding genes, followed by ncRNAs, which are also very abundant (**Main Figure 4A; Top left**). The largest functional category is receptors, followed by RNA-binding proteins, cytoskeleton, and various enzymes (**Main Figure 4A; Top Right**). Based on WormBase data, 31% of the genes identified in this study show unique muscle tissue expression (**Main Figure 4B; Top Left**). Almost all genes are present throughout development (**Main Figure 4B; Top Right**). Most genes show cytoplasmic localization, and 36% of them are expressed in operons (**Main Figure 4B; Bottom Panels**).

**Main Figure 4:**
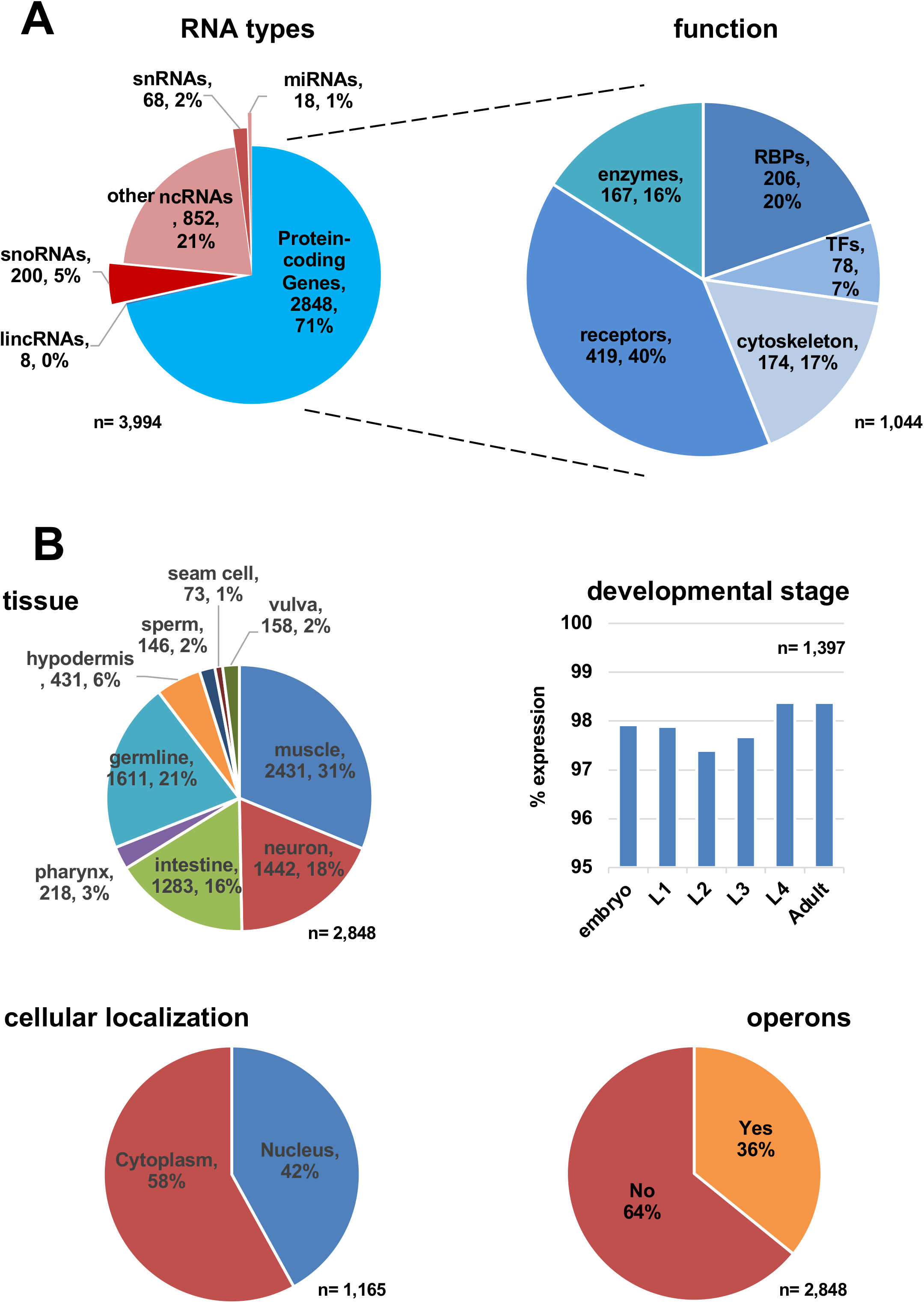
Comparative Data Analysis. A) Left: Pie chart showing the breakout of the RNA types identified in this study. Right: Out of 2,848 proteincoding genes, 1,044 genes have been previously assigned to specific biological functions and shown as a pie-chart. B) Top Left: Pie-chart showing the tissue localization of the genes identified in this study. Top Right: gene expression profile in different *C. elegans* developmental stages. Bottom Right: cellular localization. Bottom left: the number of genes identified in this study that are in operons. The comparative data used to prepare this analysis was extracted from WormBase.

### The C. elegans body muscle PROMOTERome

Next, we sought to study the core promoters of the genes identified in this study to pinpoint potential motifs transactivated by muscle-specific transcription factors. We have extracted 500 nucleotides upstream and 100 nucleotides downstream of the transcription start site of the 2,849 genes we mapped in this tissue and plotted their nucleotide distribution (**Supplemental Figure S4**). We detected an enrichment of thymidine nucleotides just upstream of the transcription start site in these genes (**Supplemental Figure S4 Panel A**). Repeated polythymine motifs (T-blocks) have been previously reported in *C. elegans* core promoters, and their presence correlates with gene expression levels [26]. We also performed a motif enrichment analysis in these core promoters and identified several novel *cis*-acting elements (**Supplemental Figure S4 Panel B** and **Supplemental Table S2**). We mapped 12 members of the human Krueppel-like transcription factors family (KLF), which are important regulators of gene expression in vertebrate development and are involved in muscle health and disease [27], Myogenin (MYOG), a muscle-specific basic-helix-loop-helix (bHLH) transcription factor involved in muscle development, myogenesis, and repair [28], and NHLH1, a helix-loop-helix transcription factor that is involved in growth and development of a wide variety of tissues and species [29] and has been linked to muscle growth and development [30]. The complete list is shown in **Supplemental Table S2**.

### The C. elegans body muscle miRNAs

We then aimed to detect and study the body muscle-specific miRNA population identified in our study. Using stringent parameters (**see Materials and Methods**), we identified 16 miRNAs expressed in the *C. elegans* body muscle tissue (**Main Figure 5A and Supplemental Table S3**). Several of these miRNAs have also been previously described to be expressed in this tissue by other groups [31], and impairing their function either leads to defects in the muscular fiber formation or shows an uncoordinated phenotype [22, 32] (**Supplemental Table S1).** We validated the body muscle localization of some of these miRNAs using two independent approaches: a GFP-based approach using worm strain expressing this fluorochrome under the control of specific miRNA promoters (**Main Figure 5B**) and a modified RT-qPCR approach using muscle-specific miRNAs immunoprecipitated from a *C. elegans* strain expressing a GFP::ALG-1 chimera in the body muscle tissue (**Supplemental Figure S5-S6**) [23, 33].

**Main Figure 5:**
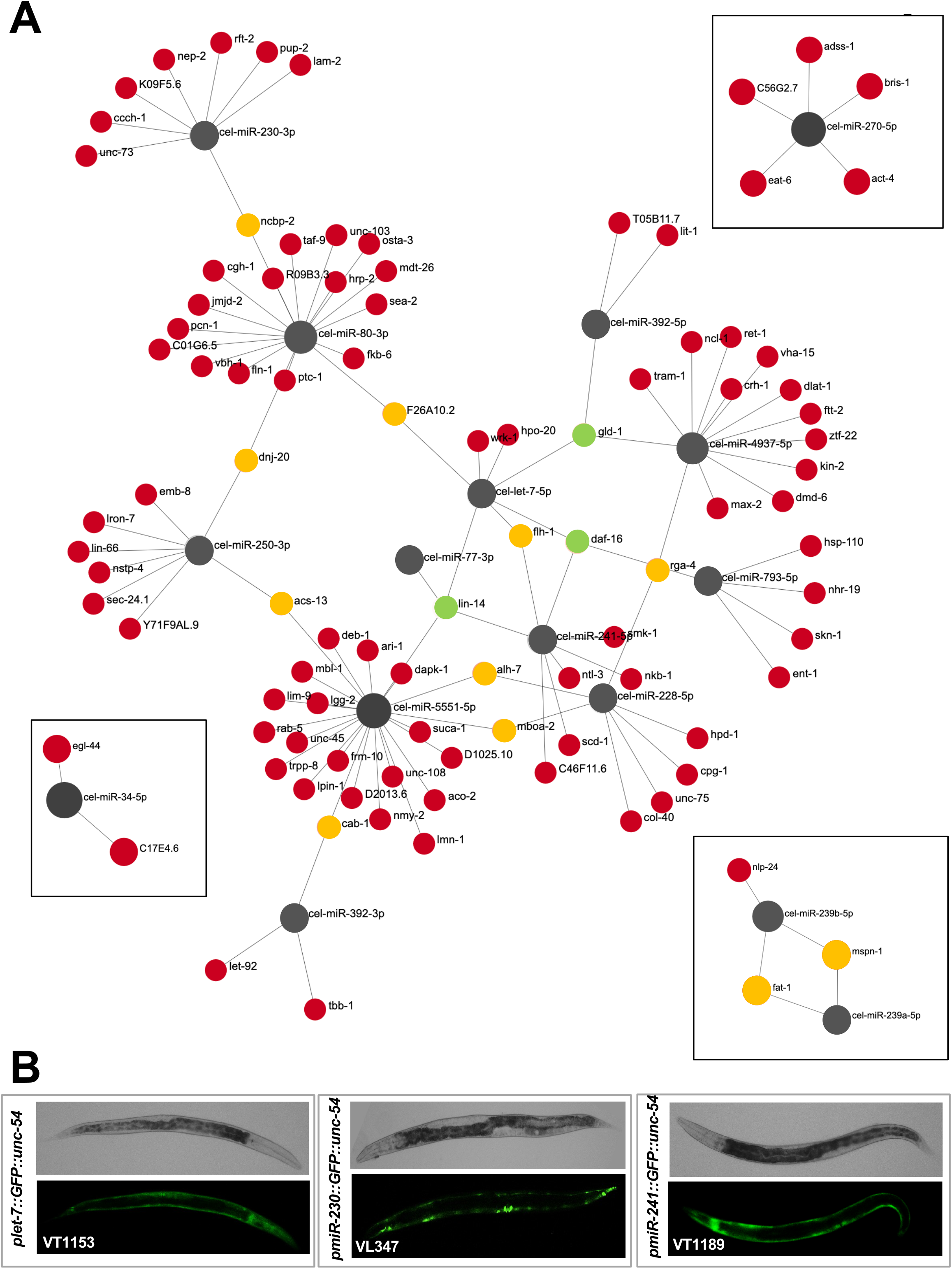
The *C. elegans* body muscle Interactome. A) The 3’UTRs of the 2,849 protein-coding genes identified in this study were screened for miRNA targets in their 3’UTRs using 16 high-quality body-muscle miRNAs identified in this study. We identified four subnetworks. Each miRNA is shown in grayscale and varies in size, depending on its number of predicted targets. Each target is shown in a different color, depending on the number of predicted miRNA targets in its 3’UTR, as from the miRanda algorithm. Red: one target, yellow: two targets, green: three or more targets. B) transgenic *C. elegans* strains expressing GFP fluorochrome under the control of three miRNAs identified in this study. *Let-7* (VT1153), *miR-230* (VL347), and *miR-241* (VT1189). All three strains show strong body muscle expression in young adult worms.

Both approaches provided strong evidence that our Nuc-Seq approach identified *bona fide* muscle-specific miRNAs. Importantly, this RT qPCR approach also allowed us to determine which one of the two miRNA strands (5p or 3p) was loaded in Argonaute. We determined that the 5p strand was preferred for *let-7, miR-34, miR-228, miR-239a, miR-239b* and *miR-5551*. The 3p strand was preferred for *miR-1, miR-77, miR-80, miR-230* and *miR-250. miR-392* did not have a distinct strand preference and was detected at lower levels than other miRNAs. We also performed RT qPCR with primers for *miR-1*, which was surprisingly below our detection threshold from our nuclear sequencing data. (**Supplemental Figures S5-S6**). With this data, we constructed a body-muscle miRNA Interactome. We performed MiRanda miRNA prediction using stringent parameters, querying the 16 miRNAs with the preferred 5p or 3p strand identified in this study to the body muscle-specific 3’UTRome data we published in the past [12, 13] network (**Main Figure 5A**).

While most body-muscle genes are targeted by a single miRNA, 11 genes are targeted by two miRNAs, two genes (*gld-1, daf-16*) are targeted by three miRNAs, and one gene (*lin-14*) is targeted by four miRNAs (**Main Figure 5A**). Our most interconnected miRNA is *miR-5551*, an uncharacterized *C. elegans*-specific miRNA predicted to target 22 muscle-expressed protein-coding genes we detected in this tissue (**Main Figure 5A**). A GO term analysis shows that its targets control contractile fibers, sarcomere, and myofibril formation (**Supplemental Figure S3**). *Let-7* and the closely related *miR-241* are also present. These miRNAs temporally regulate larval development and have been previously identified in the pharyngeal and vulval muscles [31, 34]. Among the predicted target genes, we identified several genes in the DAF insulin pathway that control muscle health and aging [35, 36].

## DISCUSSION

Here we have developed an updated FACS-based nuclear sorting method we named Nuc-Seq, to identify tissue-specific transcriptomes from cell lysates and applied it to determine the *C. elegans* muscle transcriptome. Nuc-Seq is robust, reproducible, has minimal background noise, and allows the simultaneous identification of both muscle-specific protein-coding genes and ncRNAs, such as miRNAs (**Main Figure 1**).

When applied to the *C. elegans* muscle tissue, Nuc-Seq allowed the precise identification of 2,849 protein-coding genes, including 78 muscle transcription factors and 207 non-ribosomal proteins containing an RNA-binding domain. Taken together, the muscle transcriptome we identified covers ~14% of the entire *C. elegans* protein-coding transcriptome (**Main Figure 3**). Some of these genes, such as *unc-54, mlc-1, unc-27*, and others, have been previously identified in the muscle tissue and play an important role in muscle contraction and muscle health in general. We have identified almost all genes that form the sarcomere structure and are required for muscle contraction (**Supplemental Figure S3**). When clustered using previously published protein-protein interaction data [37], the *C. elegans* muscle transcriptome is highly interconnected (**Main Figure 3B**), with several notable subnetworks corresponding to mRNA processing, protein synthesis, ATP metabolism, and muscle cell development (**Main Figure 3B**), all consistent with the function of this tissue.

Recently, similar methodologies have been utilized with success in fruit flies and mice. For example, the Fly Cell Atlas published high-quality transcriptome data from 15 somatic tissues, including other tissues that cannot be directly dissected [38]. Similarly, in mice, fluorescent nuclei have been purified from skeletal myofibers [39] and in syncytial skeletal muscle cells throughout development [40]. Together with ours, these studies highlight complex regulatory networks occurring in the muscle tissue.

We have also performed a promoter analysis, studying the promoter composition of the genes identified in this study (**Supplemental Figure S4**). Our study identified the presence of polythymine motif (T-blocks), previously reported by others, directly upstream of the transcription start site, and using prediction software (Meme Suite [41]), we have also identified many potential elements of muscle-specific transcription factors used to control gene expression in this tissue.

Our approach also allowed us to identify 16 high-quality muscle-specific miRNAs. Unfortunately, since Nuc-Seq identifies nuclear pre-miRNAs, we can only identify the miRNAs, but not which strand (5p or 3p or both) is loaded in the Argonaute protein. To bypass this limitation, we used a modified miRNA-CLIP approach coupled with RT qPCR analysis to determine which miRNA strand, the 5p or the 3p, was loaded into the Argonaute complex. We tested 11 miRNAs we identified from nuclear sequencing and additionally tested *miR-1*, which was not identified from nuclear sequencing. Interestingly, we found that *miR-1-3p* was highly expressed from our pulldown of ALG-1 in the body muscle. In contrast, some miRNAs that had high FPKM values, such as *miR-5551*, had noticeably lower expression levels from our RT qPCR data (**Supplemental Figure S6**). One critical difference between our two methods of miRNA profiling was that the miRNAs were sequenced from the nucleus instead of the cytoplasm. A possible explanation for the differences in expression is that certain miRNAs may localize to the nucleus or not be exported and associate with Argonaute proteins as mature miRNAs. Previous studies have also described nuclear-localized miRNAs [42–44] and characterized their functions [45]. Future studies investigating the miRNAs found in high abundance in our nuclear sorting but not from our cytoplasmic RT qPCR could help explain these discrepancies and if these miRNAs have biological functions.

To strengthen our miRNA Interactome results, we also ignore miRNAs described in miRbase with less than 1,000 reads. Another stringent filter we used excluded the miRNAs located within introns of other genes (mirtrons). The *C. elegans* WS250 release contains 257 miRNAs, including 118 mirtrons. We removed the mirtrons because the density of reads within their genomic regions in genes with small introns made it too difficult to map them with high confidence. This step automatically removed 46% of all *C. elegans* miRNAs from our analysis and may explain why several body muscle miRNAs identified by others [22] are not present in our list. In addition, we used very stringent filters during the miRNA identification step using the miRanda algorithm, which may have also reduced the number of target predictions.

We believe that all these stringent filters and rules, while significantly reducing the number of miRNAs and their identified target genes, increase the quality of our hits.

We have validated the body muscle localization of several miRNAs identified in our study (**Main Figure 5B and Supplemental Figures S5-S6**).

The body muscle miRNA interactome produced by clustering our 16 high-quality miRNAs and the miRanda predictions using 3’UTRs data from our 2,849 protein-coding genes identified in this study showed seven subnetworks. One large subnetwork contained 12 miRNAs (**Main Figure 5)**. 88 genes in this large subnetwork are connected to single miRNAs, but exceptions occur with *gld-1*, and *daf-16*, each targeted by three miRNAs (*let-7-5p, miR-392-5p* and *miR-4937-5p* for *gld-1*, and *let-7-5p, miR-241-5p* and *miR-793-5p* for *daf-16*). Interestingly, the heterochronic gene *lin-14* is targeted by four miRNAs in this tissue. More experiments must be performed to validate these hits and understand how and why these genes are specifically regulated in the body-muscle tissue. It this also important to note that our approach did not allow us to identify the pharyngeal muscle transcriptome, which represents the second-largest muscle tissue in *C. elegans* and could provide an important new angle to study the complete muscle tissue transcriptome and miRNAome in this organism.

In conclusion, we have produced an updated *C. elegans* body muscle transcriptome and miRNA Interactome, which will allow future studies to better understand the function of this tissue in normal states and diseases.

## MATERIALS AND METHODS

### Preparation of the transgenic strains

The *C. elegans* strain bqSi189 II; mel-28(bq5[GFP::mel-28]) III, or BN452, which ubiquitously expresses GFP and mCherry in all nuclei, was obtained from the CGC, which is funded by NIH Office of Research Infrastructure Programs (P40 OD010440). The genomic DNA was extracted from mixed stage BN452 worms, and the mCherry-tagged *his-58* sequence was cloned using PCR using BP Gateway-flanked primers specific for the Gateway entry vector pDONR221. The *mCherry::his-58* sequence was successfully cloned using Gateway technology into the entry vector, pDONR221, as evidenced by sequencing data. To assemble the finalized clone, we performed a Gateway LR Clonase II plus reaction (cat. 12538-013; Invitrogen) using the destination vector pCFJ150 (Frøkjær-Jensen *et al*. 2012) and entry clones containing the body-muscle-specific promoter *myo-3*, the *mCherry::his-58* sequence, and the *unc-54* 3’UTR as previously published (Blazie *et al*. 2017). The pCFJ150 construct and pCFJ601 (50 ng/μl), which contains a Mos1 transposon, were injected into the *C. elegans* strain EG6699 [ttTi5605 II; unc-119(ed3) III; oxEx1578] (Frøkjær-Jensen et al. 2012), which is designed for MosI-mediated single-copy integration (MosSCI), using standard injection techniques.

### Nuclei extraction

To obtain a mixed-stage population, we synchronized a population by bleaching gravid adult *C. elegans* as described previously [46]. We allowed worms to reproduce several times over approximately two weeks, which resulted in a population containing every developmental stage. Plates were inspected before nuclei isolation to confirm all life stages were present in approximately equal ratios. Six plates (100×15mm) were washed per replicate. Several medium plates of mixed stage worms were washed four times at 1,500 rpm for 3 min and resuspended in 10 mL of ice-cold NPB buffer (10□mM HEPES pH 7.6, 10□mM KCl, 1.5□mM MgCl2, 1□mM EGTA, 0.25 mM sucrose) per 250 μl volume of worm pellet. Nuclei were released by douncing 8-12 times in a chilled Wheaton stainless-steel tissue grinder (clearance 0.0005 inches =□12.5□μm) in 5 mL batches. The homogenate was sequentially filtered through 40 μm, 20 μm, and 10 μm nylon filters (pluriStrainer), then centrifuged for 10 min at 2,500 *g* at 4 °C to pellet the nuclei. The supernatant was aspirated, and the pellets containing nuclei were resuspended in 4 volumes of ice-cold NPB buffer. To remove larger debris, the resuspended nuclei pellets were centrifuged at 300 *g* for 1 min, then the supernatant containing the nuclei was transferred to a new tube on ice.

### Nuclei stability

The nuclei were isolated from the *myo-3::mCherry::his-58::unc-54* strain as described above. Nuclei were imaged in triplicates using DAPI and mCherry filters using a Leica DMi8 inverted microscope over eight hours. Images were obtained using 1 s exposure times. The result of this experiment is shown in **Supplemental Figure S1**.

### FACS sorting

Two replicates of resuspended nuclei samples for the body muscle tissue were FACS isolated into TRIzol solution using a BD Biosciences FACSAria III cell sorter with a 70 μm nozzle, and a 560 nm laser with a temperature-controlled tube holder at 4°C. The Eppendorf tubes were filled until a 1:1 TRIzol reagent and sorted nuclei prep were achieved after periodic gentle inversion. We set the gating parameters with 10,000 events from our positive control BN452 and negative control N2 and 30,000 events for the *myo-3::mCherry::his-58::unc-54* strain. We sorted 239,000 and 353,000 nuclei for the sample and the replicate for the set1 and 340,000 and 353,000 sorted nuclei for the sample and the replicate of the set2.

### RNA extraction and library preparation

The RNA from FACS isolated nuclei was extracted using an RNA MiniPrep kit (ZR2070, Zymo Research) per the manufacturer’s protocol. RNA was quantified by using a bioanalyzer. The cDNA libraries for the two replicates were generated using Nextera XT RNA sequencing. We prepared six mixed-stages cDNA libraries from the following worm strains: BN452 (two replicates), *myo-3::mCherry::his-58::unc-54* (four replicates). The libraries were depleted of ribosomal RNA contaminants using the Low Input RiboMinus Eukaryotic System v2 (ThermoFisher Scientific Catalog # A15027). The libraries were prepared from the ribosomal RNA depleted samples using the Nextflex Rapid Directional RNAseq Kit (PerkinElmer Catalog # NOVIA-5138-08) as per the manufacturer’s protocol and sequenced on the Nextseq 500 instrument (Illumina, San Diego, CA), using 75bp SE chemistry. The sequencing runs were performed at Girihlet Inc. Oakland (CA). We obtained between 50M to 60M of mappable reads across all six datasets. ***Bioinformatics analysis:*** The FASTQ files corresponding to the six datasets and the controls (total of six datasets) were mapped to the *C. elegans* gene model WS250 using the Bowtie 2 algorithm [47] with the following parameters: --local -D 20 -R 3 -L 11 -N 1 - p 40 --gbar 1 -mp 3. The mapped reads were converted into a bam format and sorted using SAMtools software using standard parameters [48]. We processed ~285M reads from all our datasets combined and obtained a median mapping success of ~93%. We used the Cufflinks software [49] to estimate the expression levels of the genes obtained in each dataset as per their BAM files. We calculated the fragment per kilobase per million bases (FPKM) number in each experiment and performed the comparison using the Cuffdiff algorithm [49]. We have pooled the four experimental samples into two sets, named set1 and set2, and the two replicates using the BN452 into a new set named ‘BN452 nuclear’. We used the median FPKM value >=5 in the set1 and set2 dataset as a threshold to define positive gene expression levels. The results are shown in **Supplemental Fig. S2 and Supplemental Table S1** using scores obtained by the Cuffdiff algorithm [49] and plotted using the CummeRbund package.

### miRNA target identification

The predicted miRNA targeting network was constructed by extracting the longest 3’UTR sequence for each of the 2,848 protein-coding genes identified in our study from the body-muscle transcriptome dataset we identified in past studies [12, 13]. We converted the 3’UTR sequences to FASTA format and parsed the file using the miRanda algorithm [50] using stringent parameters (-strict -sc −1.2). We used either the 5p or the 3p strand identified in our IP-coupled RT-qPCR approach shown in **Supplemental Figures S5-S6**. All 16 miRNAs used were identified in this study with an FPKM value >=1 (**Supplemental Table S1**). This value represents the top 5% of miRSVR scores produced by the miRanda algorithm. We also restrict the analysis to pairing scores >150 and energy score < −7. We only used miRNAs that have been previously detected in *C. elegans* with more than 1,000 reads in the miRbase database (www.mirbase.org) and that are not present in introns of protein-coding genes. The miRanda algorithm produced 118 high-quality predicted targets for 16 miRNAs. The networks were then built using the Cytoscape software [51] and uploaded to the Network Analyst online software [52] to produce the images shown in **Main Figure 5**.

### miRNA validation

To validate the RNA sequencing data shown in **Main Figure 3B**, three specific transgenic *C. elegans* strains were purchased from the CGC (VT1153, VL347, and VT1189). These strains drive GFP expression under the control of *let-7, miR-230*, or *miR-241* promoters, respectively. Each strain was imaged using identical settings for brightfield and GFP expression. These transgenic strains were imaged using a Leica DMi8 Inverted microscope. The miRNA validation in Supplemental Figure S5-S6 was performed using miRNA immunoprecipitated from a *C. elegans* strain expressing ALG-1 fused to the GFP fluorochrome expressed specifically in the body muscle tissue [23]. The immunoprecipitation step was performed using anti-GFP antibodies (Chromotek GTMA-10) as previously described [23]. Briefly, six mixed-stage 100×15mm confluent plates of transgenic *C. elegans* strains expressing the *pmyo-3::ALG-1::GFP* construct were washed 3x at 1500 rpm for 3 min. Crosslinking and RNA extraction was performed as previously described [23]. RNA was quantified, then 465ng of immunoprecipitated RNA was polyuridylated (New England Biolabs, cat no. M0337S) and reverse transcribed (Invitrogen; cat: no. 8080044) using a stem-loop poly-A reverse primer for the first strand reaction as previously described [33]. 11 miRNA-specific forward primers were designed and shown in Supplemental Table S4. Primers with low melting temperatures were not used. Polyuridynated cDNA was diluted to a 1:2 or 1:10 ratio depending on the miRNAs’ abundance. We used a 1:10 cDNA dilution for *let-7, miR-34, miR-77, miR-80, miR-228* and *miR-1* and a 1:2 cDNA dilution for *miR-230, miR-239a, miR-239b, miR-250, miR-392* and *miR-5551*. The relative abundance of each miRNA was calculated based on an exogenous positive control.

### Western Blot experiments

The western blot validation experiments shown in **Main Figure 1C** were performed using total protein for each fraction, and the input was measured using a Bradford assay; 0.75 ng of protein was used for both the nuclear and cytoplasmic fractions, and 3 ng of protein was used for the input. Primary monoclonal antibodies for RFP (6G6-100, Chromotek) (1:1000) and GAPDH (ab125247, Abcam) (1:2000) were used, followed by IRDye 800CW goat: mouse secondary antibodies (LI-COR, 925–32210) (1:5000). The membrane was imaged using the ODYSSEY CLX system (LI-COR Biosciences, NE).

### Imaging analysis

Confocal images of isolated nuclei pre-FACS stained with DAPI for the BN452 strain were acquired in the Biodesign Imaging Core, Division of the Arizona State University Bioimaging Facility. Fluorescent microscopy images of isolated nuclei pre-FACS stained with DAPI for the *myo-3::MH58::unc-54* strain were acquired using a Leica DMi8 inverted microscope.

### Network Analysis

The network shown in **Main Figure 3B** was constructed by parsing the top 2,000 hits identified in this study using the STRING algorithm (v. 11.5) [53], run with standard parameters using only ‘protein-protein interactions’ as input and set the minimum required interaction score to ‘high-confidence’ (0.700). The produced network possesses 1,995 nodes and 31,449 edges, with an average node degree of 31.5 and an average local clustering coefficient of 0.356. The network was then uploaded in Cytoscape [54] to prepare the figure.

### Promoter Analysis

We extracted 600 nt from the transcription start site for the top 100 genes identified in this study. We then used different custom Perl scripts to calculate the nucleotide distribution. The transcription factor predictions were produced by parsing these promoters to the Simple Enrichment Analysis script from the MEME suite software [41]. The results are shown in **Supplemental Figure S4** and **Supplemental Table S2**.

### Expression Analysis

The expression analysis shown in Main Figure 4 was performed using the SimpleMine tool in WormBase Field [46], which retrieves essential gene information in this database. We used as input the 2,848 protein-coding genes identified in this study. The output was then parsed with simple Perl scripts to extract and assign gene function (GO Term output), previously published tissue localization data, larval stage presence, cellular localization, and presence in operons.

## SUPPLEMENTAL MATERIALS

**Supplemental Figures S1-S6.**

**Supplemental Table S1:** Comprehensive list of transcripts identified in this study.

**Supplemental Table S2:** Simple Enrichment Analysis of sequence motifs identified in promoters of transcripts identified in this study.

**Supplemental Table S3:** miRNA target prediction of the 16 miRNAs identified in this study using the miRanda algorithm.

**Supplemental Table S4:** Primer list used in this study.

## ETHICAL APPROVAL

Not applicable.

## AUTHORS’ CONTRIBUTIONS

ALS and MM designed the experiments. ALS developed and performed the nuclear FACS sorting experiments in collaboration with AFM and MYM, and ALS prepared the sequencing reactions. MM performed the bioinformatic analysis.

ALS and MM analyzed the data, led the analysis and interpretation, assembled the figures, and wrote the manuscript. All authors read and approved the final manuscript.

## AVAILABILITY OF DATA AND MATERIALS

Raw reads were submitted to the NCBI Sequence Read Archive (http://trace.ncbi.nlm.nih.gov/Traces/sra/) with BioProject ID: PRJNA820874 and Submission ID: SUB11236444. The results of our analyses are available in Excel format as **Supplemental Table S1**.

## ACKNOWLEDGMENTS

Some of the strains used in this study were provided by the CGC, funded by the NIH Office of Research Infrastructure Programs (P40 OD010440). We thank Emma Murari and Dalton Meadows for their suggestions and advice during the preparation of the manuscript. In memory of Amelia.

