## Supplemental Material for "An updated *C. elegans* nuclear body muscle transcriptome for studies in muscle formation and function"

Additional File 1 for:

**This PDF includes:**

**Figs. S1-S6**

**Table of Contents**

### Additional File 1


**Fig. S1:** Nuclei stability overtime ……..………………….………………………………….3

**Fig. S2:** The *C. elegans* body muscle dataset .............................………….........….4

**Fig. S3:** Muscle fiber diagram showing the main components of the sarcomere .....5

**Fig. S4:** Promoter region analysis for genes detected in this study ..…………............6

**Fig. S5:** : RT qPCR approach to validate body muscle miRNAs identified from nuclear sequencing ..……………………………………………………………...............7

**Fig. S6:** miRNA validation using a modified RT-qPCR approach..……….....................8

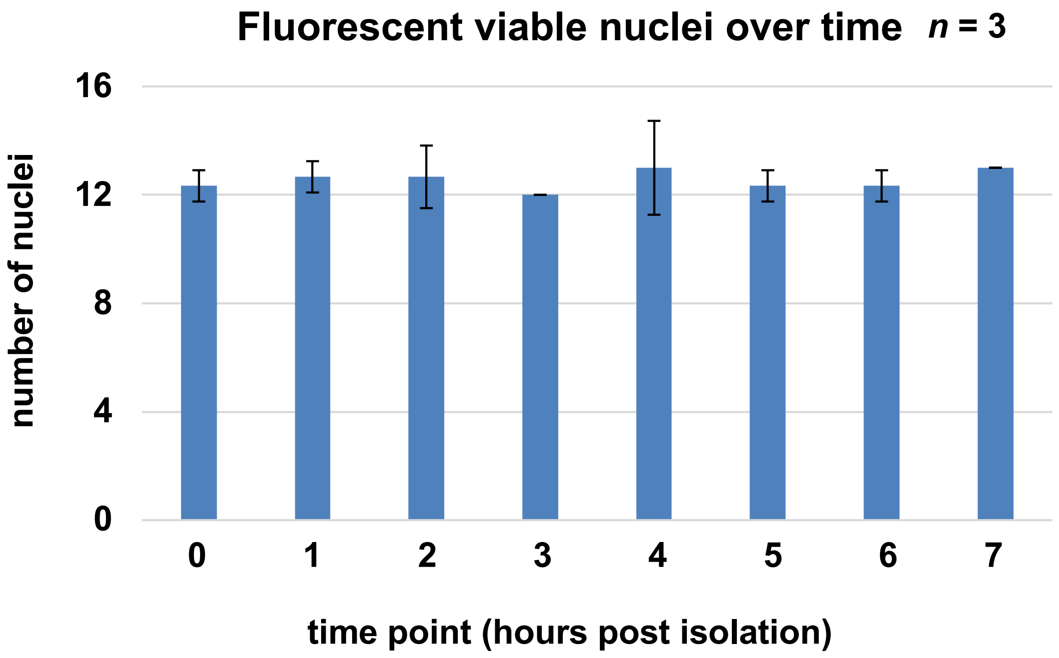


**Supplemental Figure S1: nuclei stability overtime:** The numbers of fluorescent, viable nuclei were recorded over eight time periods: time 1 as the start time, or time 0 after nuclei isolation (pre-FACS) and each time point afterwards equal to one hour on the X axis. The recorded number of mCherry fluorescent nuclei are indicated on the Y axis. Processing of the images and calculations of number of nuclei was done through imageJ software. Samples were imaged in triplicate.

**A**

| samples | | reads | | |
|---|---|---|---|---|
| | | total | mapped | mapped (%) |
| set1 | sample | 61,094,976 | 59,521,548 | 97.42 |
| | replicate | 52,785,598 | 51,118,381 | 96.84 |
| | **total** | **113,880,574** | **110,639,939** | **97.15** |
| set2 | sample | 47,432,446 | 46,155,817 | 97.31 |
| | replicate | 51198034 | 49,722,949 | 97.12 |
| | **total** | **98,630,480** | **95,878,666** | **97.21** |
| BN452 nuclear | rep1 | 43,941697 | 36,768,398 | 83.48 |
| | rep2 | 31,100,828 | 26,239,776 | 84.37 |

**B**

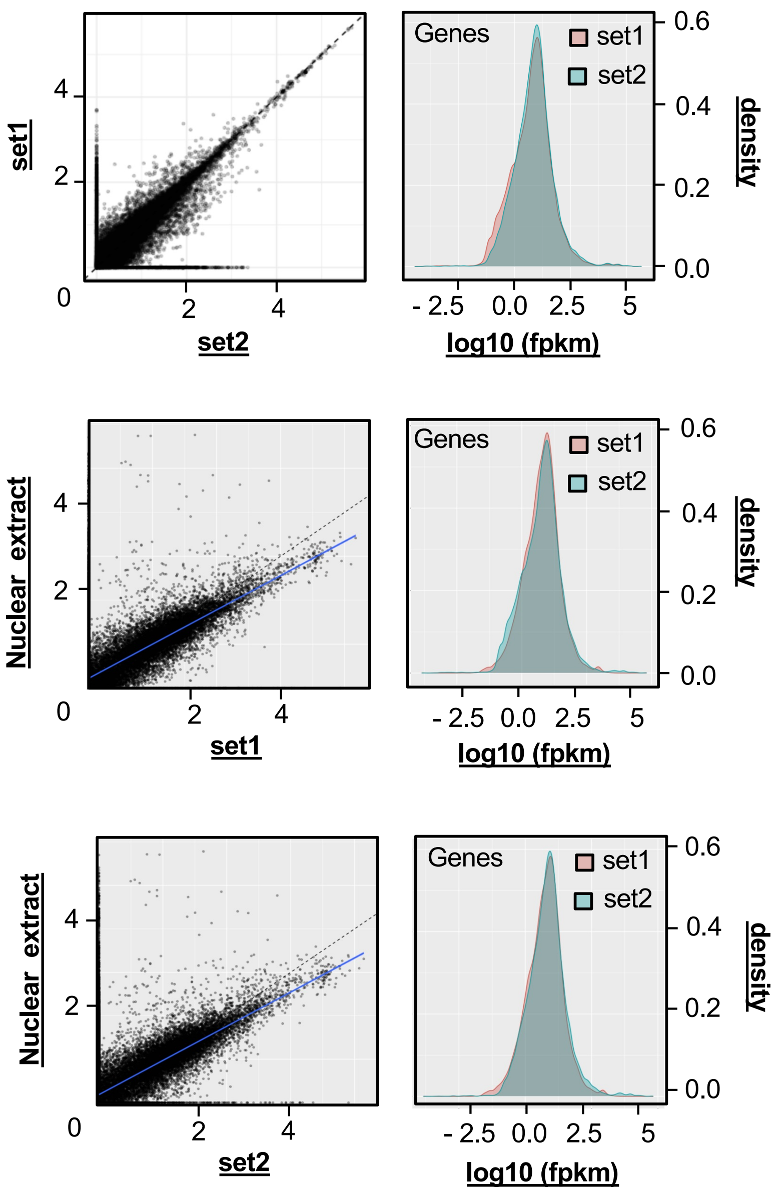


**C**

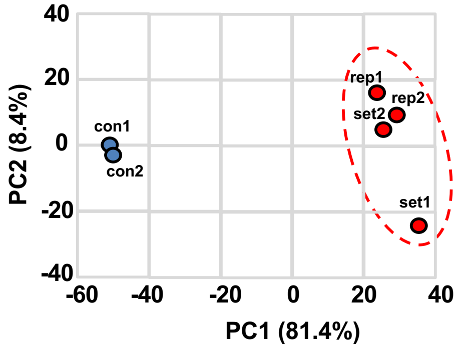


**Supplemental Figure S2: The *C. elegans* muscle Dataset**. (A) Sequencing Summary. Set1 and set2: Pooled datasets from each experimental sample and its technical replicate. 'BN452 nuclear: Pooled group containing RNA extracted from the BN452 strain (two technical replicates). (B) The distribution of the *fpkm* values in the experiment (blue) and replicate (orange) samples for each dataset. The plots were generated using the cummeRbund package v. 2.0. (C) Principal Component Analysis (PCA) shows high correlation among each duplicate within our datasets.

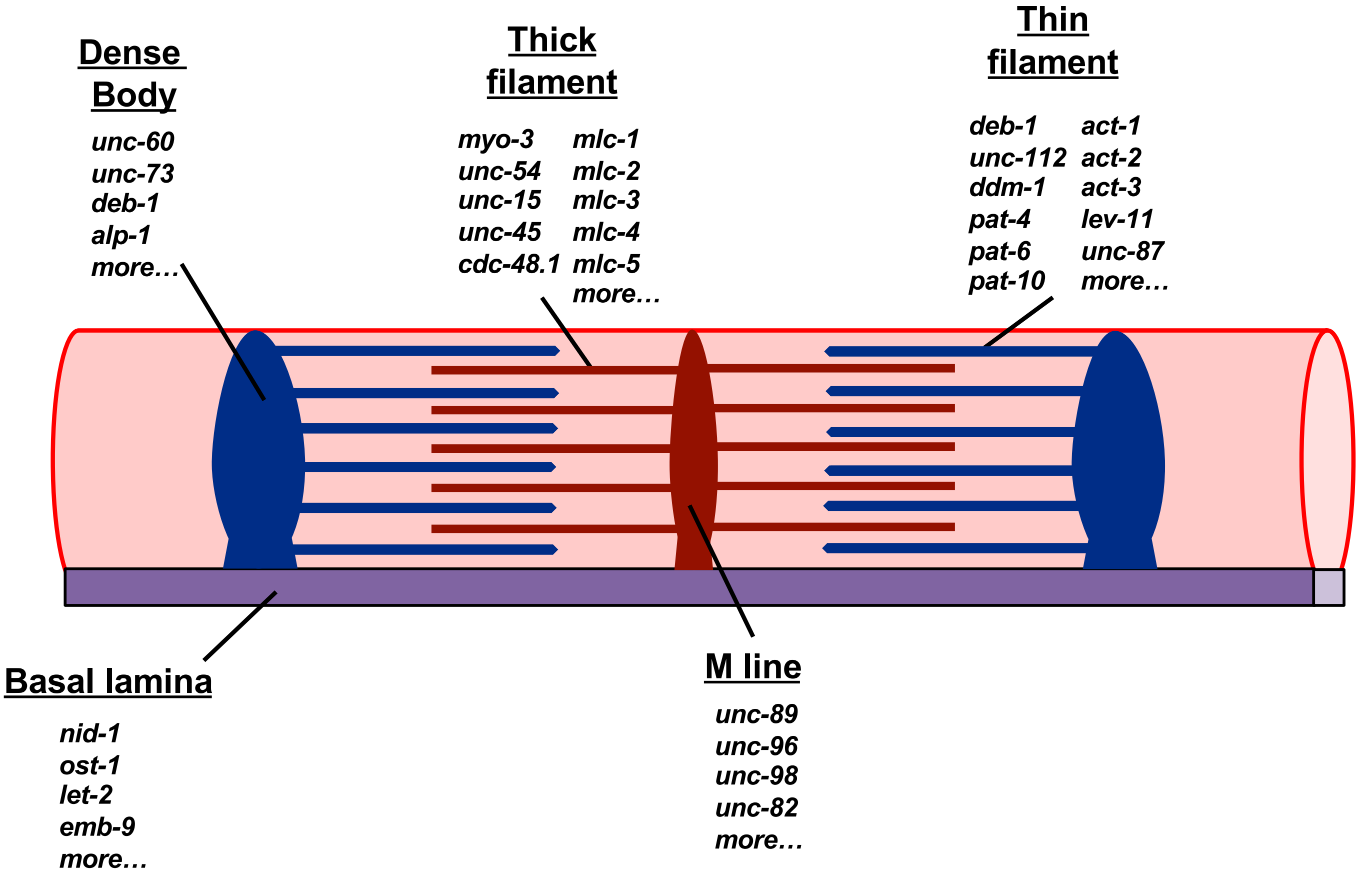


**Supplemental Figure S3: Muscle fiber diagram showing the main components of the sarcomere**. The names of the main structures are underscored. The genes shown below each structure were identified in this study and have been previously assigned as members of this structure.

# A

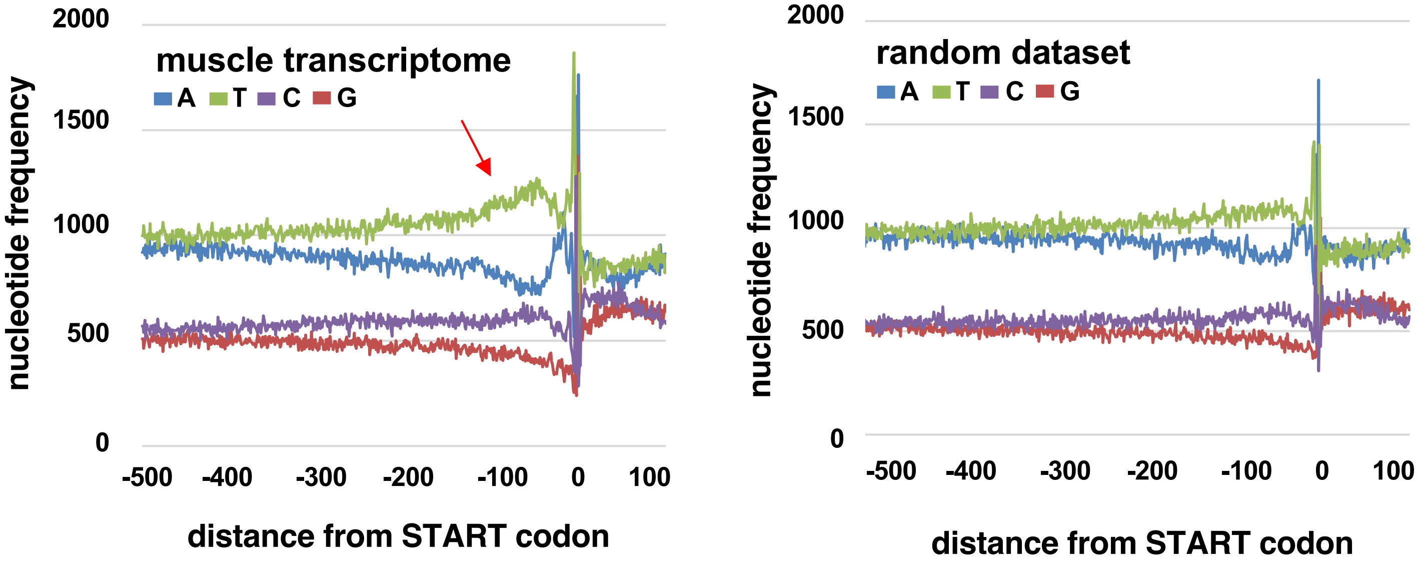


# B

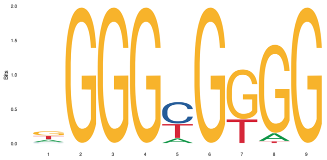

**KLF1**

| P-value | E-value | Q-value | Enrichment | Threshold |
|---|---|---|---|---|
| 9.51e-34 | 1.97e-30 | 7.77e-32 | 2.44 | 9.09 |

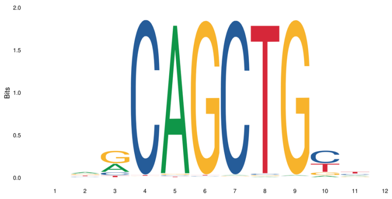

**MYOG**

| P-value | E-value | Q-value | Enrichment | Threshold |
|---|---|---|---|---|
| 4.21e-18 | 8.70e-15 | 8.34e-17 | 1.41 | 6.32 |

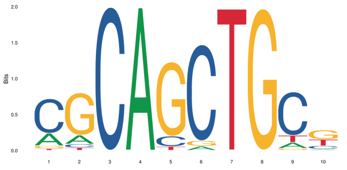

**NHLH1**

| P-value | E-value | Q-value | Enrichment | Threshold |
|---|---|---|---|---|
| 5.37e-9 | 1.11e-5 | 3.36e-8 | 1.54 | 8.80 |

**Figure S4: Promoter region analysis for genes detected in this study**. We extracted and studied the DNA regions 500bp upstream from the start codon for each of 2,848 genes unique in our muscle dataset. (A) The average base composition for promoter regions in all muscle transcripts (top), and a random dataset of the same number of genes (bottom). We detected a strong enrichment of thymidine within 100nts upstream of the transcription start site (red arrow). (B) Motif analysis. Three enriched promoter sequences are shown.

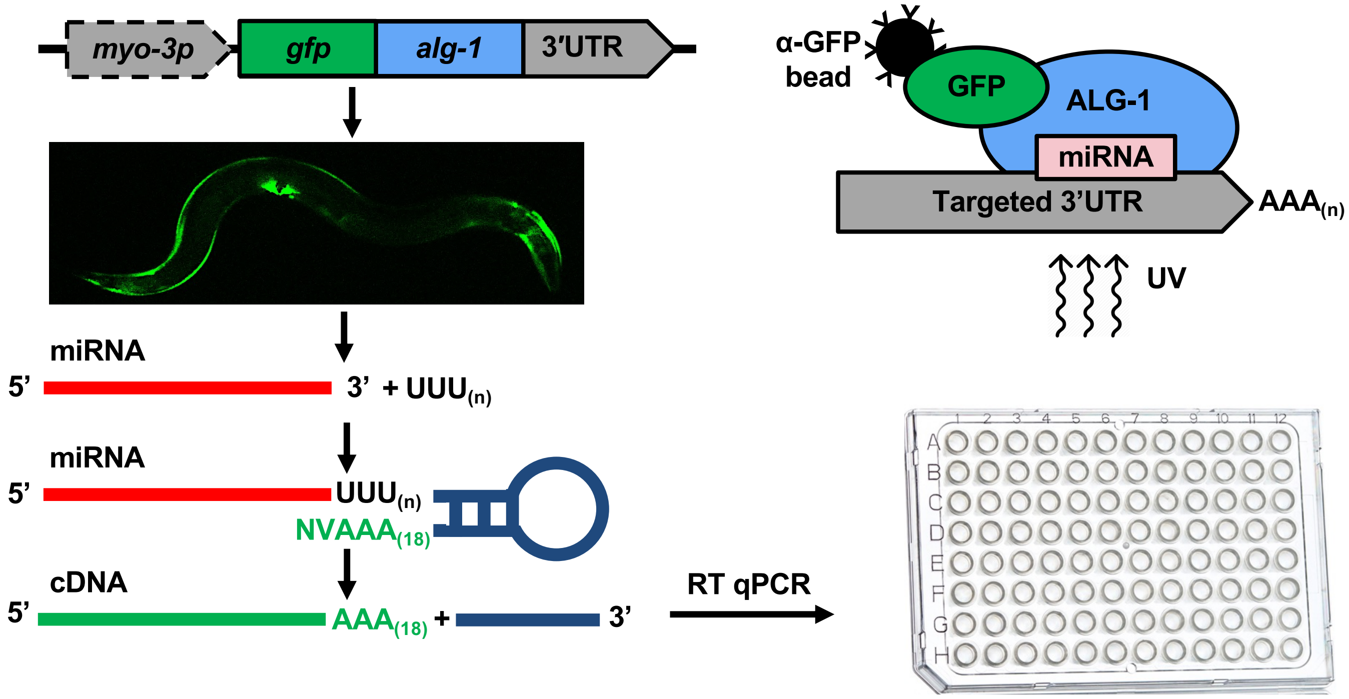


**Figure S5: RT qPCR approach to validate body muscle miRNAs identified from nuclear sequencing.** We adapted a previously described RT qPCR protocol for quantifying miRNAs. RNA was isolated from an immunoprecipitation of GFP::ALG-1 from transgenic worm strain *myo-3p::GFP::ALG-1* [23-33]. RNA was polyuridylated before performing the reverse transcriptase step using a stem-loop poly-A reverse primer containing a site for a universal reverse primer. MiRNA-specific forward primers for 5p and 3p isoforms were used to determine which strand is loaded into the Argonaute protein ALG-1.

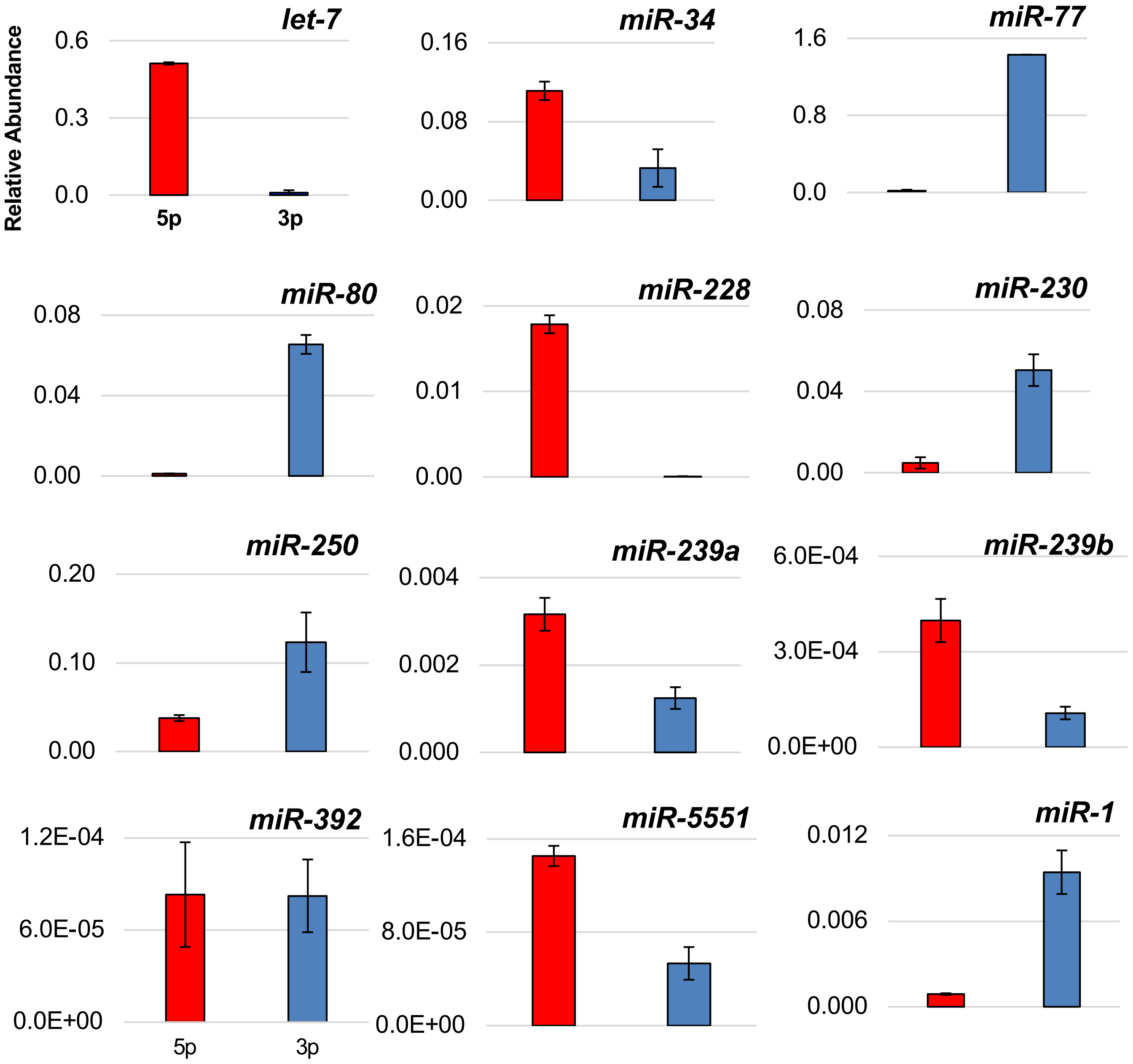


**Figure S6: miRNA validation using a modified RT-qPCR approach.** RT qPCRs were done in triplicate with forward primers specific 5p or 3p strands from miRNAs identified from body muscle tissue to demonstrate which strand is preferred for the Argonaute complex to incorporate as a mature miRNA. The relative abundance for each pair of miRNAs is shown on the Y-axis.
